# Cytokinin regulates energy utilization in *Botrytis cinerea*

**DOI:** 10.1101/2021.09.29.462332

**Authors:** Gautam Anand, Rupali Gupta, Maya Bar

**Affiliations:** Department of Plant Pathology and Weed Research, ARO, Volcani Institute, Rishon LeZion 7505101, Israel

**Keywords:** *Botrytis cinerea*, Cytokinin, Pathogenesis, Nutrition, Glycolysis, Redox

## Abstract

The plant hormone cytokinin (CK) is an important developmental regulator, promoting morphogenesis and delaying senescence. Previous work by us and others has demonstrated that CKs also mediate plant immunity and disease resistance. Some phytopathogens have been reported to secrete CKs, and may manipulate CK signaling to regulate the host cell cycle and nutrient allocation, to improve their pathogenic abilities. In a recent work, we demonstrated that CK directly inhibits the growth, development, and virulence of fungal phytopathogens, by down regulating the cell cycle and reducing cytoskeleton organization and cellular trafficking in the fungus. Here, focusing on *Botrytis cinerea (Bc)*, we report that the effect of CK on *Bc* is tied to nutrient availability; CK strongly inhibits *Bc* growth and de-regulated cytoskeleton organization in a nutrient rich environment, but has a diminished effect when nutrients are scarce. Using biochemical assays and transgenic redox sensitive botrytis lines, we examined the effect of CK on energy consumption in the fungus, and demonstrate that CK promotes glycolysis and energy consumption in *Bc*, both *in vitro* and *in planta*. Here, glycolysis and increased oxidation were stronger with waning nutrient availability. Transcriptomic data further supports our findings, demonstrating significant upregulation to glycolysis, oxidative phosphorylation, and sucrose metabolism, upon CK treatment. The metabolic effects of CK on the fungus likely reflect the role of plant CK during early infection by necrotrophic pathogens, which are known to have an initial, short biotrophic phase. In addition to the plant producing CK during its interaction with the pathogen for defense priming and pathogen inhibition, the pathogen may take advantage of this increased CK to boost its metabolism and energy production, in preparation for the necrotrophic phase of the infection. Thus, the role of CK in controlling senescence can be exploited by diverse phytopathogens to their advantage.

**Author summary:** Cytokinin (CK) is one of the primary plant developmental hormones, regulating many developmental processes. Several works have highlighted the involvement of CK in plant defense. We recently reported that CK can directly inhibit fungal plant pathogens. CK inhibits *Botrytis cinerea* growth by arresting the cell cycle and de-regulating cytoskeleton organization and cellular trafficking. Here, we report that CK positively regulates *B. cinerea* energy consumption, causing an increase in glycolytic rates and energy consumption. The effect of CK on *B. cinerea* was dependent on nutrient availability, with CK causing stronger increases in glycolysis and lower growth inhibition when nutrient availably was low, and weaker glycolytic increases coupled with stronger growth inhibition in a high nutrient environment. We propose that CK can be viewed as a bidirectional signaling molecule in plant pathogen interactions: CK acts as a signal to the fungus that plant tissue is present, causing it to activate sugar and energy metabolism pathways to take advantage of the available food source, while at the same time, CK is employed by the plant to inhibit the attacking pathogen.

## Introduction

Plant cytokinins (CKs) are known to be important in many aspects of plant life, including development of vasculature, differentiation of embryonic cells, seed development, maintenance of meristematic cells, growth and branching of root, shoot formation, chloroplast biogenesis, and leaf senescence [1,2]. CKs also play an important role in nutrient balance and stress responses in the plant [3,4]. They are known to influence macronutrient balance by regulating the expression of nitrate, phosphate and sulphate transporters [5–8]. In addition, roles for CKs in fungal pathogenesis have also been suggested, either in the context of the pathogens producing CKs, or in the context of the pathogen activating the CK pathway in the host plant [9–12]. Jameson, [13] suggested that to achieve pathogenesis in the host, CK-secreting fungal biotrophs or hemibiotrophs alter CK signalling to regulate the host cell cycle and nutrient allocation. For instance, germinating uredospores of *Puccinia spp*. have been shown to accumulate CK, modifying CK signalling to maintain plant cell cycle [14,15]. Plant CK levels can be modulated by the application of exogenous CKs, and several studies have found a positive effect of CK treatment in reduction of diseases caused by smut fungi, powdery mildew, and viruses [10,16–18]. More recently, CK was also found to enhance disease resistance to additional, non obligatory plant pathogens. Elevated levels of CKs were shown to increase host resistance to various pathogens in a wide range of plants [4,12,19– 21]. We recently reported that endogenous and exogenous applications of CKs induces systemic immunity in tomato, enhancing resistance against fungal pathogen *Botrytis cinerea* (*Bc*) by salicylic acid and ethylene signalling [4].

*Bc*, the causative agent of grey mould disease, is a cosmopolitan pathogen that can infect more than 1400 host plants, and causes massive losses worldwide annually [22]. *Bc* is a mostly [23,24] necrotrophic pathogen which has been widely used as a model pathogen to study various mechanism underlying plant-pathogen interactions. Due to its economic importance, *Bc* is listed in top 10 important plant fungal pathogens [25]. During pathogenesis, *Bc* induces necrosis in the host by producing various toxins such as botrydial, botcinic acid, and its derivatives [26,27] and production of reactive oxygen species (ROS), and also manipulates the host plant into generating oxidative bursts that facilitate colonization [28,29] and promote extension of macerated lesions by the induction of apoptotic cell death. Various enzymes including lytic enzymes, which are sequentially secreted by the fungus, facilitate penetration, colonization, and produce an important source of nutrients for the fungus [30].

Bc spores are primary sources of infection to plants in nature. After contacting the plant surface, the spores germinate to form short germ tubes that directly penetrate plant tissues [31]. It is known since long that germination of spores and infection through plant surfaces relies on their ability to access the nutrient supply offered by living plants [32,33]. Solomon and co-workers [34] have suggested a model that describes nutrient availability to the fungi during different phases of fungal infection to plant host. The first phase, which involves spore germination and host penetration, is based on lipolysis. The second phase, which requires invasion of plant tissues, uses glycolysis. Studies on *Tapesia yellundae, Colletotrichum lagenarium* and *Cladosporium fulvum* suggest that lipids are the main sources of energy during germination and penetration and after penetration the available plant sugars become the main source of energy [35–37]. These different stages of fungal infection and metabolism depend on nutrient availability and allocation. Role of CK in nutrient balance in plant is well studied but its role in nutrient balance and metabolism in necrotrophic fungus like *Bc* needs to be investigated.

Recently, we have found that CK inhibits the growth of a variety of plant pathogenic fungi. In depth characterization of the phenomenon in botrytis revealed that CK in plant physiological concentrations can inhibit sporulation, spore germination, and virulence [38] of *Bc*. We also found CK to affect both budding and fission yeast. Transcriptome profiling of *Bc* grown with CK revealed that DNA replication and the cell cycle, cytoskeleton integrity, and endocytosis, are all repressed by CK [38].

Given that CK had such fundamental, conserved, and ubiquitous effects on fungal development, the question of a possible role for CK in affecting fungal metabolism and nutrition, in particular during host-pathogen interactions, arises. In this study, we investigated the effect of CK on fungal metabolism. Using *Bc*, we examined how CK affects fungal metabolism and nutrition, both in the context of fungal growth and during infection in the tomato host. We found that CK promoted fungal metabolism, with inverse correlation to nutrient and sugar availability, and that the redox sate of *Bc* was affected by CK both during growth and during infection in the plant host. Our results reveal additional roles for CK in fungus-plant interactions, and may shed light on the availability of energy and nutrients to the fungus during initial stages of plant infection.

## Results

### CK mediated *Bc* growth inhibition and cytoskeleton de-regulation depend on nutrient availability

In order to examine if direct effect of CK on *Bc* is affected by nutrient availability, we grew *Bc* on different strengths of PDA media, with and without CK. The inhibition of mycelial growth by CK was found to depend on the media strength, and was strongest in rich media, slowly declining with decrease in nutrient availability (Fig. 1). CK-mediated growth inhibition was no longer significant in 1/8 media (Fig. 1). Similar results were obtained with growth in liquid media (S1 Fig.). Growth inhibition of *Bc* by CK in rich media was previously reported by our group [38].

**Fig. 1.**
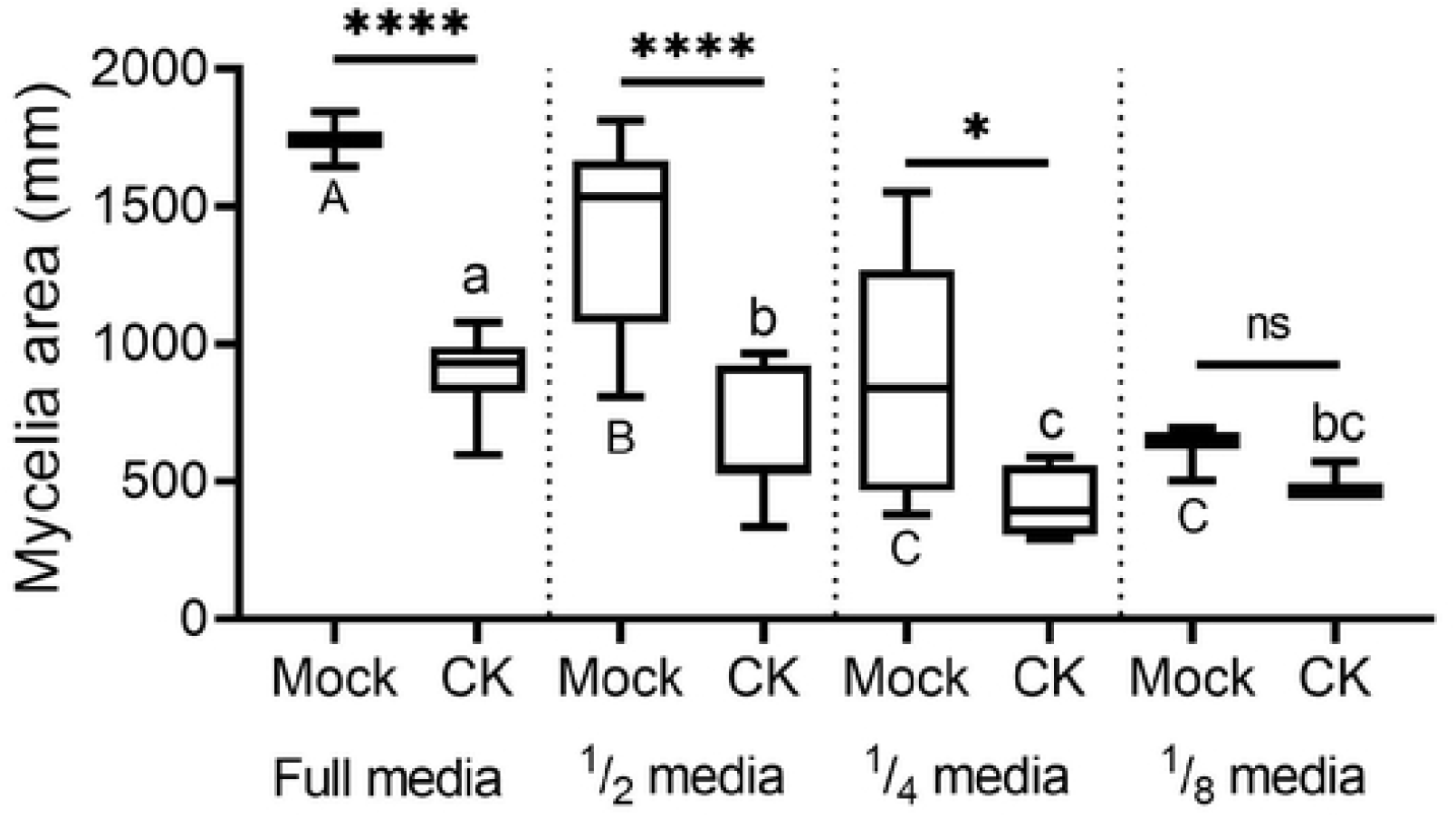
CK-mediated growth inhibition depends on nutrient availability. *Bc* mycelia were grown on PDA plates without (Mock) or with the addition of the CK 6-BAP (6-Benzylaminopurine, 100 µM) and incubated at 22 ±2 °c in the dark. Mycelial area was measured after 5 days. Boxplots are shown with minimum to maximum values, inner quartile ranges (box), median (line in box), and outer quartile ranges (whiskers), N=6. Results were analyzed for statistical significance using a one-way ANOVA with a Bonferroni post-hoc test, or a two-tailed t-test with Welch’s correction. Asterisks indicate statistically significant differences between the Mock and CK samples within the same media, *\*\*\*\*p*<0.0001; *\*p*<0.05; ns=non-significant. Upper case letters indicate statistically significant differences in the growth of Mock samples in different media, *p*<0.0035; lower case letters indicate statistically significant differences in the growth of CK-treated samples in different media, *p*<0.05.

To examine cytoskeleton integrity, we transformed *B. cinerea* with lifeact-GFP [40], and proceeded to treat the transformed fungal cells with CK in full and ¼ PDB (Fig. 2). As reported previously by our group [38], we observed mis-localization of actin, which is normally localized to growing hyphal tips [52,53], upon CK treatment in full PDB. However, there was less effect of CK on F-actin distribution when cells were grown in ¼ media (Fig. 2). Tip-specific localization of F-actin was less affected by CK in ¼ media (Fig. 2). Analysis of corrected total fluorescence in Mock and CK treated cells grown in full and ¼ PDB demonstrated that the ratio between actin in the tip of the cell, and the total cell, decreased greatly in the presence of CK in full PDB but far less in ¼ PDB (Fig. 2). An important observation was reduced actin polarization in mock samples of quarter media which might relate to reduced growth in low strength media.

**Fig. 2.**
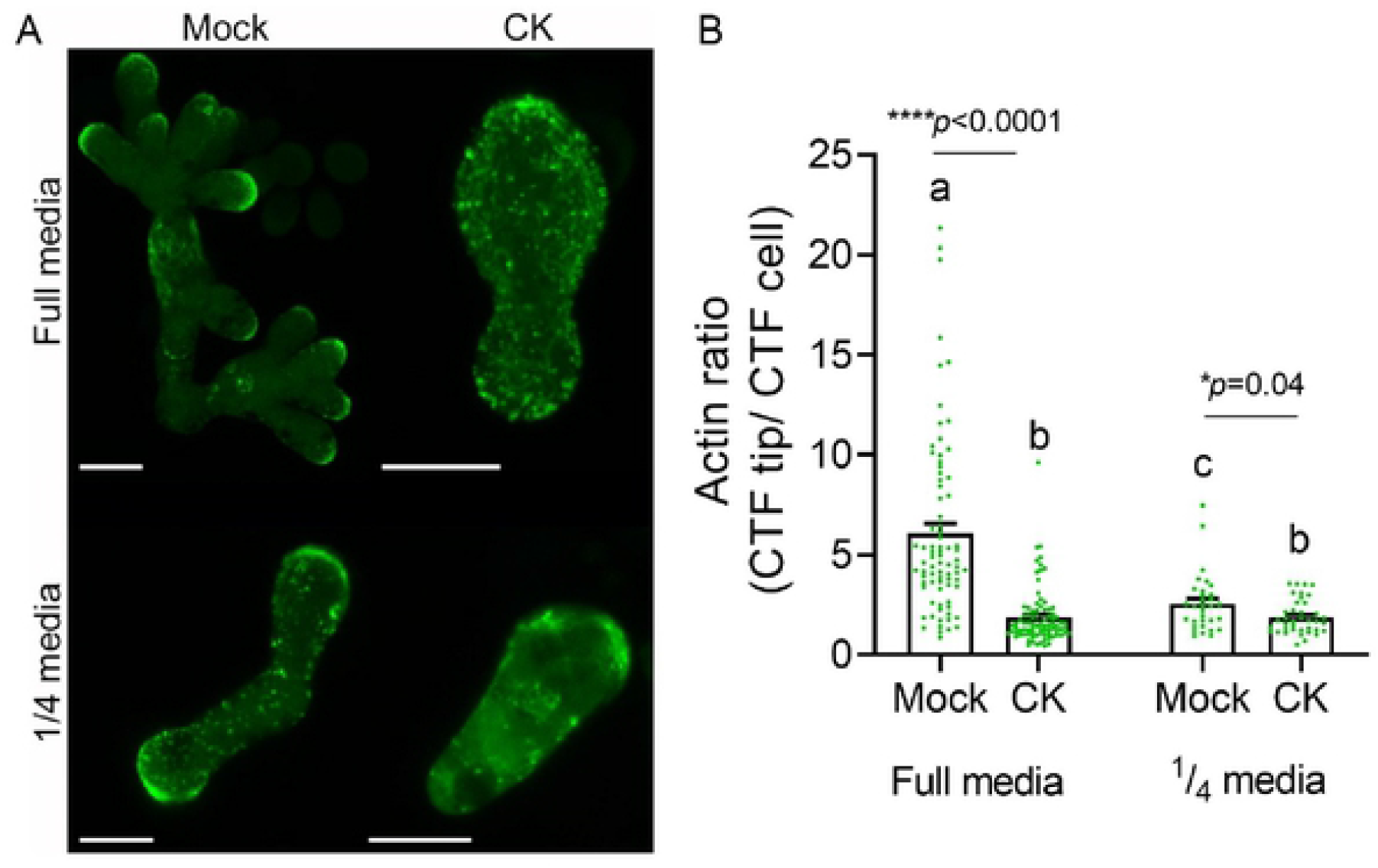
CK-mediated cytoskeleton inhibition depends on nutrient availability. Spores of *B. cinerea* expressing the filamentous actin marker lifeact-GFP, were treated with Mock or CK, and grown for 6 h and 24h hours prior to confocal visualization, in full and one forth potato dextrose broth media, respectively. **(A)** Representative images, bar=10 µM. **(B)** Analysis of corrected total fluorescence (CTF) of the ratio between actin at the tip of the cell and the total cell in Mock and CK treated cells. Three independent experiments were conducted with a minimum of 24 images analyzed, N>32 growing hypha tips. Bars are shown ±SEM, with all points. Letters and Asterisks indicate significance in Kruskal-Wallis ANOVA with Dunn’s post hoc test, \**p*<0.05 and \*\*\*\**p*<0.0001.

### Transcriptome profiling reveals an effect of CK on *Bc* metabolic pathways and sugar transport

We previously conducted transcriptome profiling on *Bc* treated with CK, finding down regulation upon CK treatment of a variety of growth and developmental pathways in *Bc*, including inhibition of cell division, DNA replication, endocytosis and the actin cytoskeleton [38]. Given that we found the effect of CK to depend on the nutritional context (Fig. 1), we next mined our transcriptomic data for alterations in gene expression that might explain this phenomenon. Interestingly, we found that the glycolysis, sucrose metabolism, and oxidative phosphorylation KEGG pathways were significantly upregulated in *Bc* upon CK treatment ([38]; Fig. 3, S1 Data). Over a quarter of the pathway genes were upregulated in the glycolysis (Figure 3A) and oxidative phosphorylation (Fig. 3C) pathways, with an FDR corrected *p-val<*0.0071 for glycolysis, and *p-val<*2.94^E-11^ for oxidative phosphorylation. A third of the sucrose/starch metabolism pathway was upregulated, FDR corrected *p-val<*0.0035 (Fig. 3D, S1 Data). Interestingly, though we found virulence genes as a group to be downregulated upon CK treatment [38], sugar-metabolism genes known to have a role in virulence such as pectin methyl esterase and poly-endogalacturonase [54], were upregulated upon CK treatment, despite the overall downregulation of virulence (Fig. 3D, S1 Data). Sugar transporters are known to be upregulated in *Bc* during pathogenesis [55,56]. In addition to glycolysis, sucrose metabolism, and oxidative phosphorylation, we found a significant upregulation of sugar transporter expression following CK treatment ([38]; S1 Data).

**Fig. 3.**
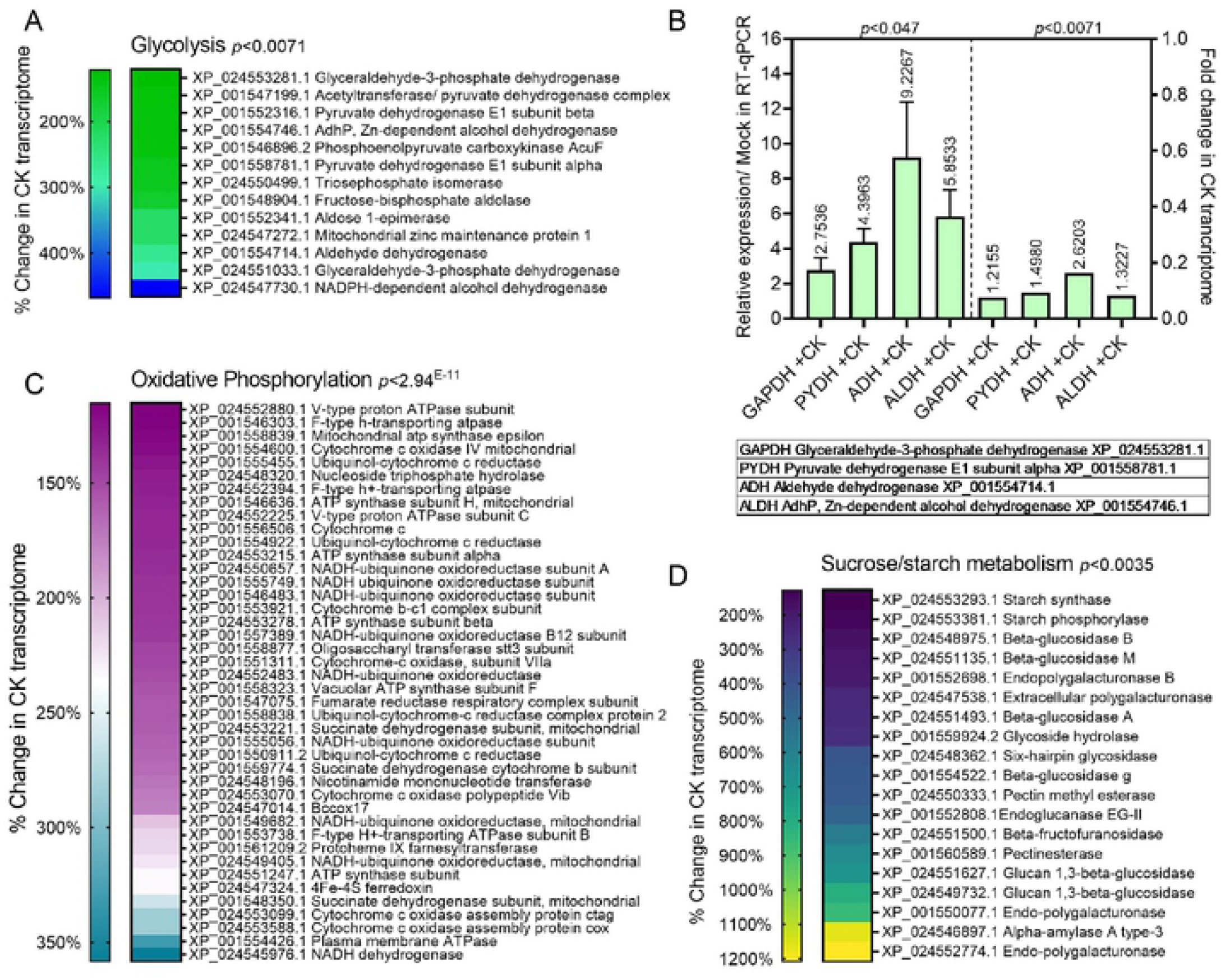
Transcriptomic analysis of botrytis grown with CK reveals up-regulation of energy metabolism pathways. Illumina Hiseq NGS was conducted on *Bc* Mock treated or CK treated samples, 3 biological repeats each. Gene expression values were computed as FPKM, and differential expression analysis was completed using the DESeq2 R package. Genes with an adjusted *p*-value of no more than 0.0S and log_2_FC (Fold Change) greater than 1 or lesser than -1 were considered differentially expressed. The KOBAS 3.0 tool was used to detect the statistical enrichment of differential expression genes in Kyoto Encyclopedia of Genes and Genomes (KEGG) pathways and Gene Ontology (GO). Pathways were tested for significant enrichment using Fisher’s exact test, with Benjamini and Hochberg FDR correction. Corrected *p*-value was deemed significant at *p<0*.*05*. The Glycolysis **(A-B)**, Oxidative phosphorylation **(C)** and Sucrose metabolism **(D)** pathways were all found to be significantly up-regulated upon CK treatment. See also Supplemental data 1. **A**,**C**,**D** Heatmap representation of upregulated genes in the CK transcriptome in each indicated pathway. **B** Comparison of RT-qPCR validation of the 4 indicated key glycolysis genes with the transcriptomic values. The full transcriptome data was previously published (Gupta et al., 2021) and is available (NCBI bioproject PRJNA718329).

For glycolysis pathway genes, we also conducted a RT-qPCR validation of the upregulation of some of the key pathway genes found to be upregulated in the transcriptome following CK treatment. Fig. 3B depicts a comparison between the fold change of these genes in the transcriptome, and the changes we observed in an independent experiment in qPCR.

### CK rescues inhibition of glycolysis and ATP synthesis in a nutrient availability dependent manner

To examine to effect of CK on *Bc* metabolism, we used 2-Deoxy-D-glucose (2-DG) and oligomycin (OM), which are inhibitors of glycolysis and ATP synthesis, respectively. 2-DG is a glucose analog that inhibits glycolysis by competing with glucose as a substrate for hexokinase, the rate-limiting enzyme in glycolysis. After entering the cell, 2-DG is phosphorylated by hexokinase II to 2-deoxy-d-glucose-6-phosphate (2-DG-6-P) but, unlike glucose, 2-DG-6-P cannot be further metabolized by phosphoglucose isomerase. This leads to the accumulation of 2-DG-P in the cell, and subsequent depletion in cellular ATP [57]. OM inhibits mitochondrial H^+^-ATP-synthase, and has been attributed antifungal properties [58,59]. *Bc* was grown with these two inhibitors separately at different media strength, as described above. 2-DG significantly inhibited growth at low media strength, in the concentration used (Fig. 4). OM was inhibitory at all media strengths in the concentration used (Fig. 4). CK (100 µM) was found to rescue the inhibitory effect of 2-DG when added to the growth media. Lowering the nutrient availability promoted CK-mediated rescue of glycolysis inhibition by 2-DG. Interestingly, following OM treatment, the fungi became insensitive to CK and were not further inhibited by the addition of CK. A partial rescue of ATP synthesis mediated inhibition of growth by CK was observed in full and 1/2 media. These results strengthen the notion that CK is affecting glycolysis in the fungus, and is in the same pathway as ATP synthesis.

**Fig. 4.**
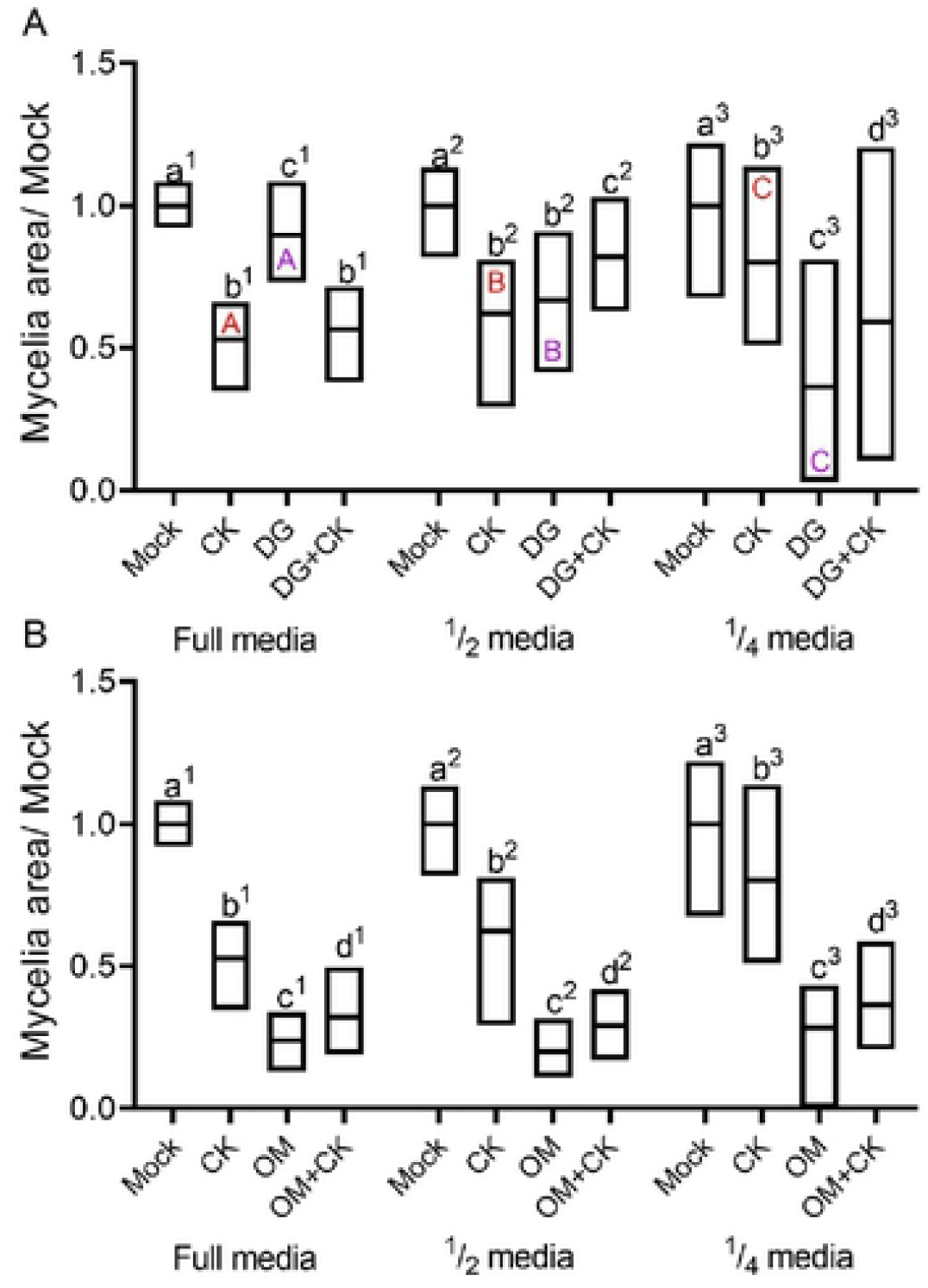
CK rescues glycolysis inhibition and partially rescues ATP synthesis inhibition. *Bc* mycelia were grown on PDA plates without (Mock) or with the addition of the CK 6-BAP (6-Benzylaminopurine, 100 µM), the competitive glucose inhibitor 2-DG (2-deoxyglucose, 2.5 mM) (**A**), or the ATP synthesis inhibitor OM (oligomycin, 1 µM) (**B**) and incubated at 22 ± 2 °C in the dark. Mycelia area was measured after 5 days. Floating bars are shown with minimum maximum values, line in bar indicates median. ±SEM, N=10. **A**: 2-deoxyglucose (DG). Results were analyzed for statistical significance using a one-way ANOVA with a Tukey post-hoc test. Lower case letters indicate statistically significant differences between samples, with number tags indicating the group that was comparatively analyzed, *p*<0.025. Upper case letters within the top of CK bars indicate statistically significant differences in the level off CK-mediated growth inhibition, *p*<0.018. Upper case letters within the bottom of DG bars indicate statistically significant differences in the level off DG-mediated growth inhibition, *p*<0.011. **B**: Oligomycin (OM). Results were analyzed for statistical significance using a one-way ANOVA with a Tukey post-hoc test, or a two-tailed t-test with Welch’s correction. Letters indicate statistically significant differences between samples, with tags indicating the group that was comparatively analyzed, *p*<0.038.

### CK induces glucose uptake in *Bc*

The dependence of CK-mediated growth inhibition on nutrition and energy state, and the rescue of glycolysis inhibition by CK, indicated that CK is affecting *Bc* metabolism. To further confirm this hypothesis, we next measured glucose uptake in the presence of CK in both rich PDB and synthetic defined liquid media. We observed a significant increase of glucose uptake in the presence of CK in both media types (Fig. 5). Interestingly, the percent increase of glucose uptake by *Bc* in the presence of CK increased with decreasing nutrient availability in the media (Fig. 5). There was significant increase of sugar uptake in defined media but it was not dependent on media strength. We know 2-DG competes with glucose in the glycolytic pathway. The increase of sugar uptake might be the reason why CK rescues glycolysis inhibition by 2-DG. Since ATP is required for growth, it is not surprising that inhibition of ATP synthesis prevented further growth inhibition by CK.

**Fig. 5.**
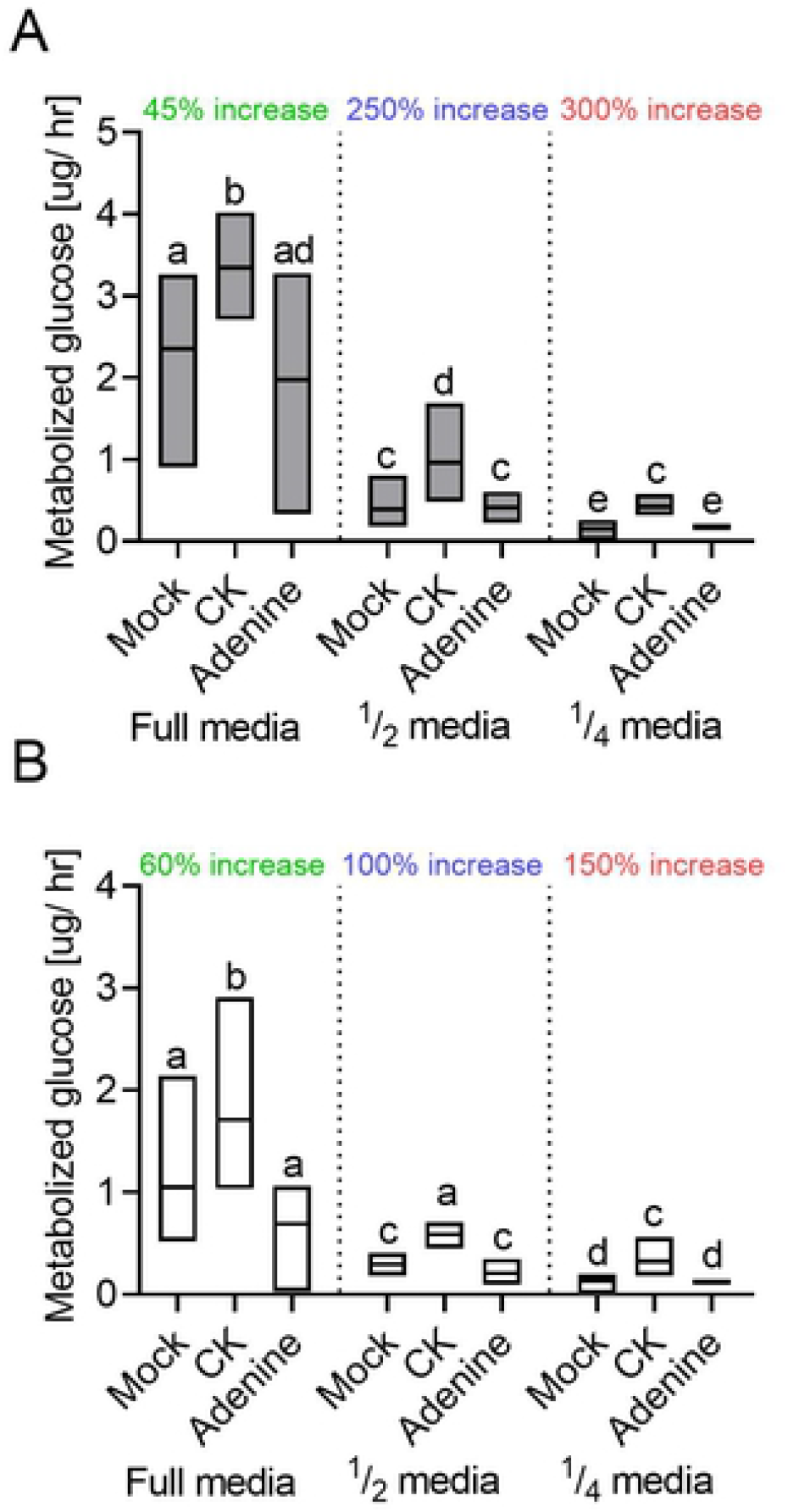
CK promotes glucose uptake. *Bc* spores (10^6^/ ml dissolved in sterile water) were grown in PDB (**A**) or Synthetic medium (**B**) with 150 rpm shaking, at 22 ± 2 °C in the dark, without (Mock) or with the addition of the CK 6-BAP (6-Benzylaminopurine, 100 µM), or the structural control Adenine, 100 µM. The amount of glucose in the media was examined after 48 h, and subtracted from the amount of glucose present in media without fungi that underwent similar treatment. The approximate percent of increase in glucose uptake in the presence of CK is indicated above the bars for each media concentration. Floating bars are shown with minimum maximum values, line in bar indicates median. N=6. Results were analyzed for statistical significance using two-tailed t-test with Welch’s correction. Letters indicate statistically significant differences between samples, **A** *p*<0.04, **B** *p*<0.049.

### CK alters *Bc* redox status

Since metabolic pathway fluctuations can affect redox status [60], and redox homeostasis in *Bc* is known to change during host infection [61,62], we next examined cytosolic and mitochondrial redox status in the fungus grown with CK. For this purpose, we generated *Bc*l-16 strain lines expressing GRX-roGFP and mito-roGFP, using previously described expression cassettes that were used for the measurement of *Bc* redox status [40,51]. We found that after 24 hours of growth, CK significantly altered the cytosolic redox of *Bc* to a more reduced state, while the mitochondrial redox was significantly more oxidised with CK (Fig. 6A). Following redox over time, we observed similar states in the first 8 hours of growth, with CK starting to affect the redox state after about 15 hours of co-cultivation, corresponding with the stage at which mycelia are elongating (Fig. 6B-C). Cytosolic and mitochondrial redox states are often inversely correlated [50]. Interestingly, there was an inverse effect of CK on cytosolic and mitochondrial redox of growing mycelia (Fig. 6).

**Fig. 6.**
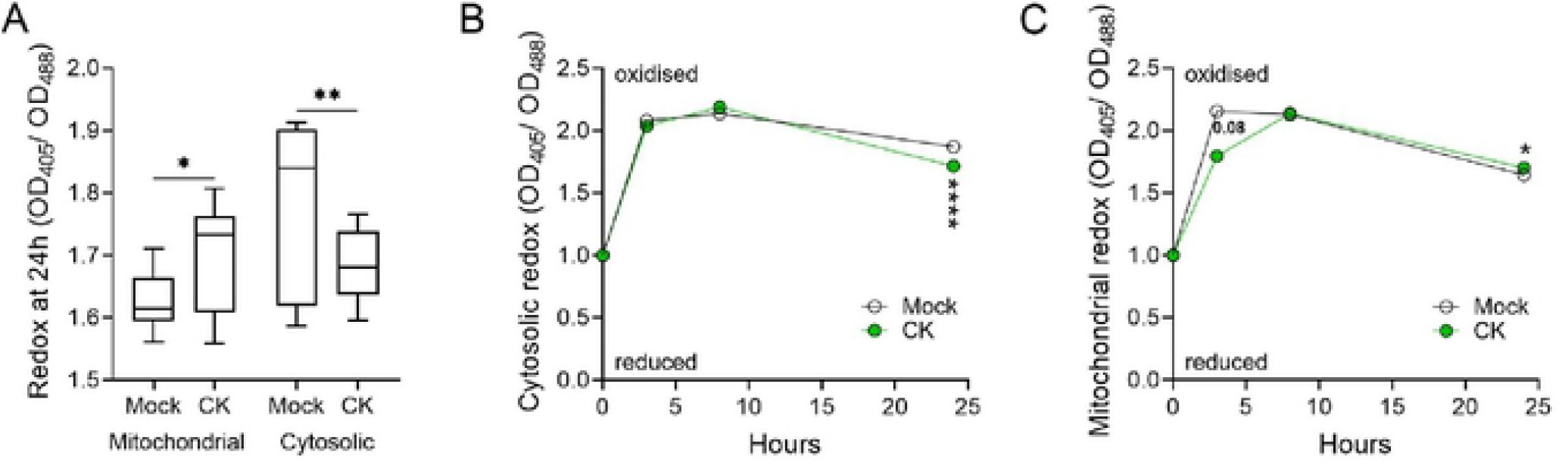
CK alters *Bc* redox state in rich media. The redox state of *Bc* without (Mock), or in the presence of CK, was assessed using roGFP transformed *Bc*. Spores (10^6^/ ml) of *Bc* strains expressing GRX-roGFP, for assessing cytosolic redox, and mito-roGFP, for assessing mitochondrial redox, were incubated in PDB without (Mock) or with CK 6-BAP (6-Benzylaminopurine, 100 µM), for 24 h at l8 °C, with 150 rpm shaking. Fluorescence was measured using a fluorimeter, with excitation at 405 ± 5 nm for the oxidized state and 488 ± 5 nm for the reduced state of roGFP2. The emission was detected at 510 ± 5 nm. The redox ratio of the fungus was calculated as Em405/Em488 of Relative fluorescence units (RFU). (**A**) Redox status of the mitochondria and cytosol, with and without CK, after 24 h. Boxplots are shown with minimum to maximum values, inner quartile ranges (box), median (line in box), and outer quartile ranges (whiskers), N=6. (**B**) Time course of redox state in the cytosol. (**C**) Time course of redox state in the mitochondria. Asterisks indicate statistical significance in a two-tailed t-test, *\*p*<0.05, *\*\*p*<0.01, *\*\*\*\*p*<0.0001.

### Endogenous CK content of tomato leaves affects redox state of *Bc* during plant infection

Since CK affected redox in *Bc* in rich media, we next examined whether endogenous CK content in tomato leaves can affect the redox status of infecting *Bc* mycelia. For this, *Bc* GRX-roGFP and mito-roGFP conidia from freshly sporulated PDA plates were used to infect detached leaves from M82, IPT, and CKX plants. Leaves overexpressing IPT have increased CK content and are more resistant to *Bc* infection, while leaves overexpressing CKX have reduced CK content and are more sensitive to *Bc* infection [4]. Redox-dependent changes in GRX-roGFP2 and mito-roGFP fluorescence in living Botrytis hyphae have been previously visualized by confocal laser scanning microscopy (CLSM) [50]. Infecting *Bc* hyphae expressing GRX-roGFP in the cytosol or mito-roGFP in the intermembrane mitochondrial space were analysed microscopically, 24 and 48 hours after inoculation. Similar to the fluorometry-based calculations, a higher 405nm/488nm ratio indicates more oxidised state, and a lower ratio, a more reduced state. We found that 24 hours after inoculation, the cytosolic redox state of the infecting hyphae was more oxidised on IPT leaves (high CK content) as compared to the infecting hyphae on mock M82 leaves, while infecting hyphae on CKX (low CK content) were more reduced (Fig. 7A,C). In a parallel set of experiments which included additional genotypes, we also observed increased oxidation of the *Bc* cytosol when infecting M82 leaves that were pre-treated with CK, or when infecting leaves of the hypersensitive *clausa* mutant (S2 Fig.). 48 hours post inoculation, we observed an opposite trend of the cytosolic redox of the infecting hyphae, with hyphae on IPT becoming more reduced while hyphae on CKX were more oxidised, when compared with M82 leaves. (Fig. 7A,C). Here, again, in a parallel set of experiments, which included additional genotypes, we also observed increased reduction of the *Bc* cytosol when infecting leaves of the hypersensitive *clausa* mutant (S2 Fig.). When examining mitochondrial redox of the hyphae growing on IPT or CKX leaves, we found that, 48 post inoculation, mitochondrial redox of the hyphae growing on IPT was significantly oxidised as compared to mock M82, while hyphae growing on CKX were significantly reduced (Fig. 7B,C). The cytosolic redox was measured in a parallel set of experiments on leaf discs co-cultivated with *Bc* spores in a fluorimeter plate, with similar results (S3 Fig.). To verify that the virulence of the roGFP fungi was intact, we also conducted disease assays of these fungi on the different genotypes, with findings consistent with previous results [4], i.e., reduced disease on high-CK or CK-hypersensitive genotypes, and increased disease on low-CK genotypes (S4 Fig.).

**Fig. 7.**
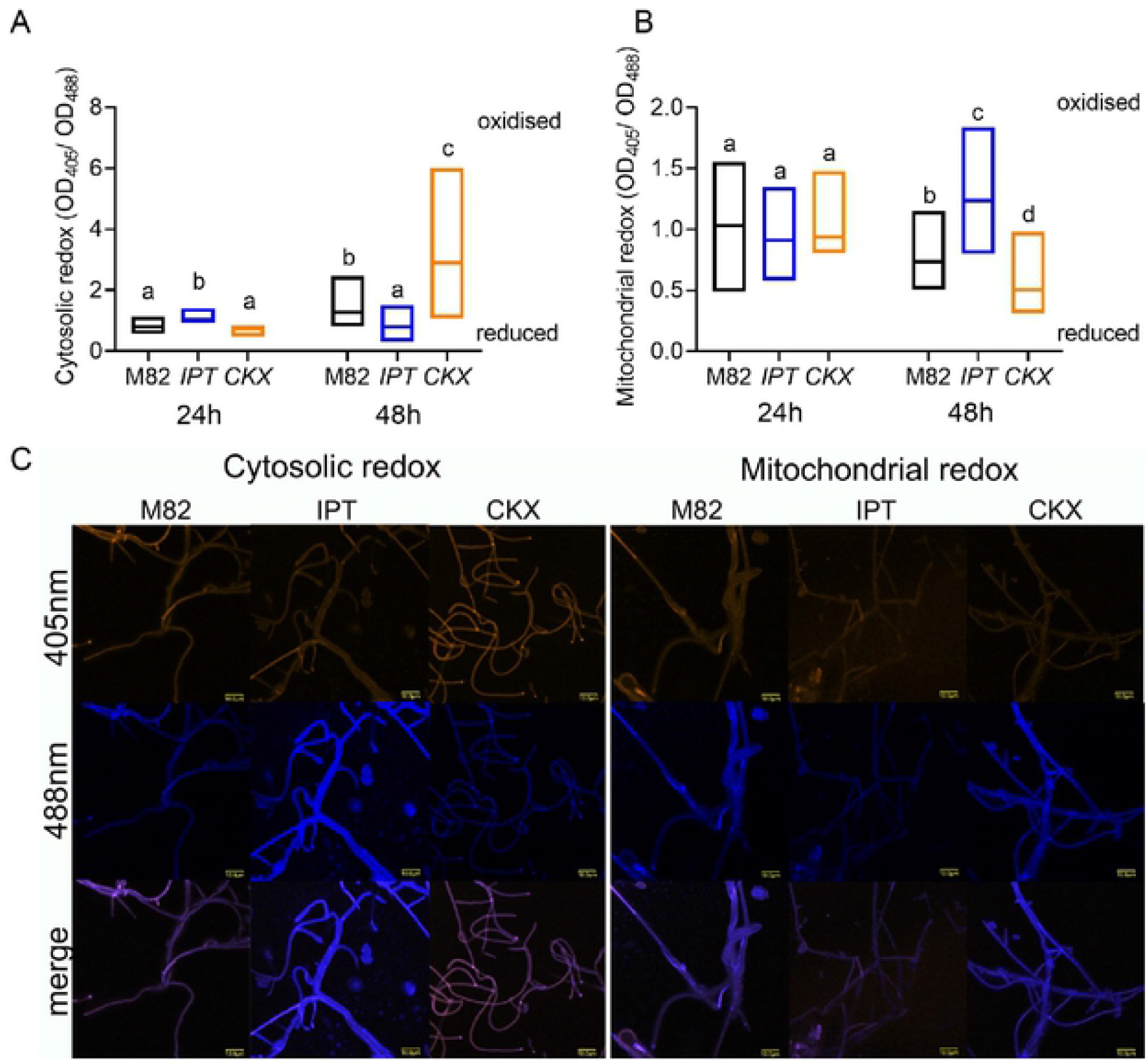
Plant endogenous CK alters *Bc* redox state during infection. The redox state of *Bc* when infecting leaves of different CK-content tomato genotypes was assessed using roGFP transformed *Bc*. Spores (10^6^/mL in glucose and K_2_HP0_4_) of *Bc* strains expressing GRX-roGFP, for assessing cytosolic redox, and mito-roGFP, for assessing mitochondrial redox, were used to infect the background M82 wild-type line, the high-CK *pBLS>>IPT7* overexpressing line (“IPT”), and the low-CK *pFIL>>CKX3* overexpressing line (“CKX”). *Bc* fluorescence was captured using a confocal laser scanning microscope at 24 h and 48 h, with excitation at 405 nm for the oxidized state and 488 nm for the reduced state of roGFP2. The emission was detected using a 505-530 nm bandpass filter. The redox ratio of the fungus was calculated as Em405/Em488 using lmageJ, from at least 12 images per time point, per treatment. (**A**) Redox status of the *Bc* cytosol, after 24 h and 48 h. (**B**) Redox status of the *Bc* mitochondria, after 24 h and 48 h. (**A-B**) Floating bars are shown with minimum to maximum values, lines indicates median, N=12. Differences between samples were assessed using a one-way ANOVA with a Dunnett post hoc test. Different letters indicate statistically significant differences between samples, (A)*p*<0.021, (B)p<0.029. (**C**) Representative images of the roGFP fungi growing on leaves of the different genotypes, captured at the “reduced” and “oxidized” wavelengths.wavelengths.

We further examined the transcriptome of *Bc* grown in the presence of tobacco seedlings following CK treatment, finding significant changes, both significant downregulation and significant upregulation, in NADPH/NADH reductases and oxidoreductases (S5 Fig.). These transcriptional changes in the fungus further support the notion that CK is affecting ROS coping mechanisms in *Bc*, and could underlie the altered pathogenesis courses observed in tomato genotypes with altered CK content.

## Discussion

It has been previously reported by us and others that CK promotes fungal disease resistance in plants [4,12]. Recently, we reported a direct inhibitory effect of CK on *Bc* growth and development *in vitro* [38]. Our previous results indicated that *B. cinerea* responds to CK and activates signaling cascades in its presence, leading to inhibition of the cell cycle, mis-localization of the actin cytoskeleton, and inhibition of cellular trafficking [38]. Our previous RNAseq data provided several clues as to which pathways in the fungus are affected by CK. Given those results, in addition to previous observations concerning the cell cycle and cytoskeleton of the fungus in the presence of CK, we hypothesized that CK may be exerting its effect through influence on fungal metabolic pathways. The present study was performed to examine the effect of CK on *Bc* metabolism. We exmined sugar uptake, glycolysis, and cellular redox status of *Bc* in the presence of CK. Our results demonstrate that the inhibitory activity of CK against *Bc* is largely dependent on the status of nutrient and energy availability. We found that the inhibitory effect of CK on *Bc* was correlated with nutrient and energy availability, with fungi grown in rich media being inhibited more strongly than fungi grown in sub-optimal conditions (Fig. 1, S1 Fig.). Intact F-actin was found to be required for hyphal growth, morphogenesis, and virulence, which were all impaired in F-actin capping protein deletion mutants [63]. Since we had previously observed that CK caused mis-localization of actin at the growing tip of *Bc* hyphae [38], we examined whether this phenomenon was also correlated with the nutritional status of the environment. Indeed, we found that the effect of CK on the cytoskeleton is also dependent on the status of nutrient availability (Fig. 2). Interestingly, we observed reduced polarization of F-actin in *Bc* growing in minimal media when compared with rich media (Fig. 2), providing one underlying mechanism for reduced fungal growth under low nutrient conditions.

Our previously published transcriptome profile revealed that important developmental pathways such as cell division, cellular trafficking, and the cytoskeleton, were inhibited upon CK treatment. Our results demonstrated that the effect of CK on *Bc* is dependent on nutritional status. Re-examining our transcriptomic data in light of this, we found that glycolysis, sucrose metabolism and oxidative phosphorylation pathways were significantly enriched upon CK treatment (Fig. 3). The expression of sugar transporters was also significantly upregulated in the RNAseq data.

This transcriptomic data was generated under defined nutrient conditions. To further examine possible effects of CK on fungal metabolism, we investigated the effect of CK on *Bc* glycolysis and ATP synthesis, under different nutrient and energy availability. We found that CK was able to rescue inhibition of glycolysis and ATP synthesis in a nutrient dependent manner, with stronger rescue observed under minimal nutrient conditions. CK also promoted an increase in sugar uptake by the fungus, in a nutrient dependent manner, with the strongest uptake promotion observed under minimal nutrient and energy conditions (Figs. 4-5). Upregulation of glycolysis and oxidative phosphorylation key genes, together with the increased uptake of sugar, could explain the rescue of metabolic inhibition by CK. Taken together, these results confirm that the effect of CK on *Bc* is dependent on nutrient availability.

Changes in metabolic pathways are often reflected in the redox status [62]. Hence, we examined the effect of CK on *Bc* redox status both *in vitro* and *in planta*, using tomato genotypes with varying CK content or sensitivity. *Bc* cells grown with CK in rich media had a significantly reduced cytosol and a significantly oxidized mitochondria (Fig. 6). A reduced cytosol and oxidized mitochondria is indicative of increased glycolysis and oxidative phosphorylation [64,65]. Thus, this result correlated with our transcriptomic data, in which glycolysis and oxidative phosphorylation are upregulated in the presence of CK (Fig. 3), and also with our results demonstrating that CK promotes sugar uptake in *Bc* (Fig. 5). *In planta*, after 48h of inoculation, *Bc* had a reduced cytosol and oxidized mitochondria when infecting the CK-rich IPT, and an oxidized cytosol and reduced mitochondria when infecting the CK-deficient CKX, confirming that CK can affect the redox state of *Bc* during infection *in planta*. This significant change in redox was also coupled with the lower virulence on IPT leaves and higher virulence on CKX leaves (S4 Fig., [4]). It was previously reported that the cytosol of infecting *Bc* hyphae on dead onion peel is reduced [50]. Our results also show that *Bc* cytosol is more reduced in IPT, but the resultant infection was lower when compared to that on CKX. In addition to the different hosts systems and different infection time frames, a possible explanation for this could be the CK-mediated immunity induced in the host [4,12]. The results of redox state of *Bc* hyphae in vitro and in planta, together with transcriptome data and sugar uptake results, might relate to the role of CK in nutrient allocation in fungi during infection.

CKs have previously described roles in plant–pathogen interactions [3,4]. The interaction of some biotrophic pathogens with their hosts leads to the formation of green bionissia (formerly known as green islands) [66], which are sites of green living tissue surrounding the sites of active pathogen growth [9]. The formation of these green bionissia is correlated with elevated levels of cytokinins in these tissues. It is believed that cytokinins likely delay the onset of senescence in green bionissia, allowing pathogens access to more nutrients from the plant [9]. Necrotrophic fungal pathogens, which obtain their nutrients from dead plant cells, have also been reported to cause the formation of green tissue around the sites of infection (green necronissia) in certain cases [9]. Application of exogenous CK is known to induce the formation green necronissia [9,15,67]. We found that the effect of CK on *Bc* is dependent on nutrient availability, and can induce glycolysis, oxidative phosphorylation and sugar uptake. We also observed CK-mediated redox shifts in *Bc*, that are likely due to these increases in metabolism.

We previously observed that CK inhibits *Bc* growth and development in vitro, in plant-physiological concentrations. The metabolic effects of CK on the fungus likely reflect the role of plant CK during early infection by necrotrophic pathogens, which have been demonstrated they have a short biotrophic phase [23,68]. Necrotrophic pathogens secrete toxins and enzymes to cause cellular damage. Thus, in addition to the plant producing CK during its interaction with the pathogen for the purpose of priming its defenses and inhibiting pathogen growth, the pathogen may take advantage of this increase in CK to exploit the formation of green necronissia, by increasing its metabolism and energy production, to prepare for the necrotrophic phase of the infection. Thus, the role of CK in controlling senescence, which is used by biotrophic pathogens to their advantage, likely also benefits necrotrophs during their biotrophic phase. However, when CK content is high as in IPT, the CK is also directly inhibitory against the pathogen, causing an attenuation of virulence, rendering the advantage of the CK to the pathogen in this initial phase of the infection irrelevant.

Our work suggests that CK may serve as central player in the hormonal cross-talk between plant host and phytopathogen, and that the role of CK in controlling senescence can be exploited by diverse fungal phytopathogens to their advantage. Future research will focus on the role of CK in nutrient allocation in fungal phytopathogens during infection, affording insights into fungal infection phases in the context of host-phytopathogen interactions.

## Materials and Methods

### Fungal growth conditions

*B. cinerea* strain Bcl16 (*B. cinerea, Bc* was grown on potato dextrose agar (PDA) at 22 ± 2 °C for 5 days. Bcl-16 sporulates well on different types of media including PDA [38].

### Mycelial growth assay

To study how nutrient availability affects mycelial growth of *Bc* in the presence of CK, 6-BAP (6-Benzylaminopurine, Sigma-Aldrich) was dissolved in 10mM NaOH and added to PDA media of full, half, one-fourth and one-eighth strength. To study how *Bc* responds to metabolic inhibitors in the presence of CK, 2-Deoxy-D-glucose (2-DG) and oligomycin (OM) were added to PDA media at above mentioned strength to final concentration 100 µM, 2.5 mM and 0.1 µg/mL respectively. *Bc* mycelial plugs (5 mm) taken ∼1cm from the edge of a fresh plate were placed at the centre of PDA plates and incubated under the above mentioned growth conditions. To measure the mycelium weight in liquid media, Bc was cultured in stationary liquid PDB full and one-fourth media strength in the presence of 100 µM concentrations of 6-BAP. After 72 h, the fungal mass was dried and the dry weight was measured.

### Effect of CK on glucose uptake at different media strength

To evaluate the effect of CK on glucose uptake, spores were harvested in 1 mg mL^-1^ glucose and 1 mg mL^-1^ K_2_HPO_4_ and filtered through 40 μm pore cell strainer. Spore concentration was adjusted to 10^6^ spores mL^-1^ using a Neubauer chamber. *Bc* spores were grown in potato dextrose broth (PDB) or defined media of full, half and one-fourth strength. Composition of defined media was glucose (20 g/L) and 4 g/L each of K_2_HPO_4_, KH_2_PO_4_, and NH_4_SO_4_. 100 µM of CK were added to both PDB and defined media cultures, which were then allowed to grow for 48 hours. The amount of metabolized glucose was analysed by measuring the amount of glucose present in the media by standard DNSA method [39] using dextrose as standard. Metabolized glucose was assumed to be inversely proportional to the amount present in the media. For control, glucose in different strength of media subjected to above mentioned conditions but without *Bc* was measured.

### Generation of *B. cinerea* lines expressing lifeact-GFP

For generation of *Bc* strains expressing Lifeact-GFP, we used a fusion construct to target replacement of nitrate reductase (bcniaD) gene in manner reported previously by our group and others [38,40]. The plasmid pNDH-OLGG, which is used as a template for the amplification of expression cassette, has flanking sequences of bcniaD, a resistance cassette for hygromycin, and the filamentous actin (F-actin) imaging probe “Lifeact” fused to GFP. Primers GA 34F/34R (S1 Table) from our previous study [38] were used for the amplification of the expression cassette. PEG-mediated transformation was used to transfer the PCR amplified expression cassette to *Bc* [41]. Fungal transformants were visualized under a confocal microscope and screened with primers GA 44F/44R and GA 31F/31R (S1 Table). To examine the effect of CK on cytoskeleton in different nutrient availability, spores of transformed Bc were treated with Mock or CK (100uM) in full and ¼ PDB and grown for 6h and 24h hours respectively, prior to confocal visualization in full and one forth PDB broth media, respectively. We acquired confocal microscopy images using a Olympus IX 81 inverted laser scanning confocal microscope (Fluoview 500) equipped with an OBIS 488 nm laser lines and a 60× 1.0 NA PlanApo water immersion objective. GFP images of 24 bits and 1024 × 1024 pixels were imaged using the excitation/emission filters: BP460-480GFP/BA495-540GFP. Image analysis was conducted with Fiji-ImageJ using the raw images and the 3D object counter tool and measurement analysis tool [42].

### Transcriptome analysis of metabolic pathway genes

Procedures for RNA preparation, quality control, sequencing, and transcriptome analysis are detailed in our previous work [38]. Differential expression analysis was executed using the DESeq2 R package [43]. Genes with an adjusted *p*-value of no more than 0.05 were considered differentially expressed. PCA was calculated using the R function prcomp. The sequencing data generated in this project was previously published [38], and the raw data is available at NCBI under bioproject accession number PRJNA718329.

The gene sequences were used as a query term for a search of the NCBI non-redundant (nr) protein database that was carried out with the DIAMOND program [44]. The search results were imported into Blast2GO version 4.0 [45] for gene ontology (GO) assignments. Gene ontology enrichment analysis was carried out using Blast2GO program based on Fisher’s Exact Test with multiple testing correction of false discovery rate (FDR). KOBAS 3.0 tool (http://kobas.cbi.pku.edu.cn/kobas3/?t=1) [46] was used to detect the statistical enrichment of differential expression genes in KEGG pathway and Gene Ontology (GO).

### Effect of CK on the expression of glycolysis genes

To examine the effect of cytokinin on fungal glycolysis and validate the RNAseq results, we grew *Bc* spores in PDB with 0 and 100 µM 6-BAP in a rotary shaker at 180 rpm and 22 ± 2 °C for 24 hours. Total RNA was isolated using Tri reagent (Sigma-Aldrich) according to the manufacturer’s instructions. RNA (3μg) was used to prepare cDNA using reverse transcriptase (Promega, United States) and oligodT15. qRT-PCR was performed on a Step One Plus Real-Time PCR system (Thermo Fisher, Waltham, MA, United States) with Power SYBR Green Master Mix protocol (Life Technologies, Thermo Fisher, United States). For glycolysis analysis, we selected the following genes: Glyceraldehyde-3-phosphate dehydrogenase (XP_024553281.1), pyruvate dehydrogenase (XP_001558781.1), aldehyde dehydrogenase (XP_001554714.1), and alcohol dehydrogenase (XP_001554746.1). The primer sequences for each gene, and primer pair efficiencies, are detailed in S2 Table. A geometric mean of the expression values of the three housekeeping genes: ubiquitin-conjugating enzyme E2 (ubce) [47], Iron-transport multicopper oxidase, and Adenosine deaminase [48] was used for normalization of gene expression levels. All primer efficiencies were in the range 0.97-1.03 (S2 Table). Relative expression was calculated using the copy number method [49]. At least six independent biological replicates were used for analysis.

### Generation of redox sensitive *Bc*

For the detection of redox state in cytosol and mitochondria, *B. cinerea* expressing GRX-roGFP and mito-roGFP were generated. For the expression of the redox sensors, the construct generated previously was used [40,50,51]. For generation of constructs expressing GRX-roGFP at the bcniaA locus, we used the plasmid pNAH-GRX-roGFP as template. The vector contains 5′ and 3′ flanking sequences of bcniaA, a resistance cassette mediating resistance to hygromycin, and sensor for the redox potential of the cellular glutathione pool glutaredoxin probe “GRX” fused to GFP. The expression cassette carrying the hygromycin resistance gene, GRX-roGFP and the bcniaA flanking sequence was amplified using primers GA 41F/41R (S1 Table). The PCR amplified expression cassette was used to transform *B. cinerea* using PEG mediated transformation. 0.125% lysing enzyme from *Trichoderma harzianum* (Sigma– Aldrich, Germany) was used for protoplast generation. Following PEG mediated transformation, protoplasts were plated on SH medium containing sucrose, Tris-Cl, (NH_4_)_2_HPO_4_ and 35µg/mL hygromycin B (Sigma–Aldrich, Germany). Colonies that grew after 2 days of incubation were transferred to PDA-hygromycin medium, and conidia were spread again on selection plates to obtain a monoconidial culture. Fungal transformants were visualized under a confocal microscope and screened with primers GA 42R/42R (S1 Table). Confirmed transformants were stored at -80°C and used for further experiments.

### Measurement of *Bc* redox status in liquid media

Redox state of roGFP transformed *Bc* was measured in liquid culture using a fluorimeter. *Bc* strains expressing GRX-roGFP and mito-roGFP were grown for 2 weeks on PDA medium at 18°C in the light to induce mass sporulation. 10 mL of PDB medium containing 100 µM CK was inoculated with conidia and incubated for 24 h at 18 °C on 150 rounds per minute (rpm). 1mL samples were taken and washed twice in double distilled water. A 96-well plate (Microplate pureGrade™ 96-well PS, transparent bottom) was inoculated with 200 µL of the washed germlings and used for fluorescence measurements using a fluorimeter (Promega GloMax® explorer multimode microplate reader, GM3500, USA). Fluorescence was measured at the bottom with 3 × 3 reads per well and an excitation wavelength of 405 ± 5 nm for the oxidized state and 488 ± 5 nm for the reduced state of roGFP2. The emission was detected at 510 ± 5 nm [50]. The gain was set to 100. Relative fluorescence units (RFU) were recorded to calculate the Em405/Em488 ratio.

### Imaging of *Bc* redox state during on-plant pathogenesis

To examine the effect of endogenous CK content on the redox state of *Bc* during pathogenesis, Bc GRX-roGFP conidia from freshly sporulated PDA plates were used to infect detached leaves from the *S. lycopersicum* M82 background line, as well as M82 overexpressing the Arabidopsis isopentenyl transferase (IPT) gene *AtIPT7* under the FIL promoter: *pFIL>>IPT7* (IPT), and M82 overexpressing the Arabidopsis cytokinin oxidase (CKX) gene *AtIPT3* under the BLS promoter: *pBLS>>CKX3* (CKX) [4]. For *Bc* inoculation, *Bc* was grown on PDA in the dark at 22±2°C. Ten days old plates were given daylight for 6 h, and then returned to the dark for sporulation. Spores were harvested in 1 mg mL^-1^ glucose and 1 mg mL^-1^ K_2_HPO_4_, and filtered through 40 μm pore cell strainer. Spore concentration was adjusted to 10^6^ spores mL^-1^ after quantification under a light microscope using a Neubauer chamber. Leaflets 10-15 days old tomato plants were excised and immediately placed in humid chambers. Leaflets were inoculated with one droplets of 5 μL suspension. Twenty-four hours after inoculation, germinated conidia were imaged using a fluorescent Olympus IX 81 inverted laser scanning confocal microscope (Fluoview 500). Images were collected with a 60× 1.0 NA PlanApo water-immersion lens in multi-track line mode. roGFP was excited at 405 nm in the first track and at 488 nm in the second track. For both excitation wavelengths, roGFP fluorescence was collected with a bandpass filter of 505–530 nm and averaged from four readings for noise reduction [50]. The Ratiometric analyses of fluorescence images were calculated using Fiji-ImageJ.

### Data analysis

Data is presented as minimum to maximum values in boxplots or floating bar graphs, or as average ±SEM in bar graphs. For Gaussian distributed samples, we analyzed the statistical significance of differences between two groups using a two-tailed t-test, with additional post hoc correction where appropriate, such as Welch’s correction for t-tests between samples with unequal variances. We analyzed the statistical significance of differences among three or more groups using analysis of variance (ANOVA). Regular ANOVA was used for groups with equal variances, and Welch’s ANOVA for groups with unequal variances. Significance in differences between the means of different samples in a group of three or more samples was assessed using a post-hoc test. The Tukey post-hoc test was used for samples with equal variances, when the mean of each sample was compared to the mean of every other sample. The Bonferroni post-hoc test was used for samples with equal variances, when the mean of each sample was compared to the mean of a control sample. The Dunnett post-hoc test was used for samples with unequal variances. For samples with non-Gaussian distribution, we analyzed the statistical significance of differences between two groups using a Mann-Whitney U test, and the statistical significance of differences among three or more groups using Kruskal-Wallis ANOVA, with Dunn’s multiple comparison post-hoc test as indicated. Gaussian distribution or lack thereof was determined using the Shapiro-Wilk test for normality. Statistical analyses were conducted using Prism8^™^.

## Acknowledgments

This work was supported by the Israel Science Foundation grant No. 1759/20 to MB. The lifeact and roGFP constructs were kindly provided by Julia Schumacher. GA is supported by the Indo-China ARO Postdoctoral Fellowship Program. MB thanks members of the Bar group for continuous discussion and support.

## Author contributions

Conceptualization: GA, MB. Design: GA, RG, MB. Methodology & experimentation: GA, RG. Analysis: GA, RG, MB. Manuscript: GA, MB.

**The authors declare no competing interest**

## Data availability statement

The authors declare that the data supporting the findings of this study are available within the paper and its supplementary information files. Raw data is available from the corresponding author upon reasonable request. The raw data generated in the transcriptomic analyses is deposited in NCBI under Bioproject accession number PRJNA718329.

## Supplementary information

**S1 Fig**. CK-mediated growth inhibition depends on nutrient availability-Dry weight measured from fungi grown in liquid media.

**S2 Fig**. Plant endogenous CK alters *Bc* cytosolic redox state during infection-additional genotypes.

**S3 Fig**. Plant endogenous CK alters *Bc* redox state during infection-plate assay.

**S4 Fig**. *Bc* transformed roGFP lines display virulence behaviour similar to that of the background line.

**S5 Fig**. CK alters *Bc* redox state-changes in the transcriptome.

**S1 Table**. Oligonucleotides used for generating and validating *Botrytis cinerea* transgenic strains.

**S2 Table**. Primers used in RT-qPCR.

**S1 Data**. Transcriptomic effect of CK on *Bc* metabolic pathways.

